# Profiling Chromatin Accessibility in Humans Using Adenine Methylation and Long-Read Sequencing

**DOI:** 10.1101/2023.10.05.561129

**Authors:** Vahid Akbari, Sreeja Leelakumari, Steven J.M. Jones

**Author notes:** Correspondence (S.J.M.J.).

## Abstract

Chromatin accessibility demonstrates accessible DNA regions that usually have regulatory function. Several studies have demonstrated DNA accessibility profiling using nanopore sequencing and GpC or adenine modification. However, the GpC dinucleotide is not evenly distributed across the genome and there are regions sparse in GpC. 6mA studies have demonstrated chromatin accessibility in yeast using nanopore based sequencing, although a high-false positive in 6mA detection rate and low coverage have previously precluded the effective use in mammalian sized genomes. Here, we have developed accurate 6mA base calling and demonstrated high-resolution profiling of accessible regions and simultaneous CpG methylation detection in humans using long read sequencing.

## Introduction

Eukaryotic DNA is packaged via various proteins to form chromatin and to compactly fit in the nucleus. This packaging also results in DNA to be inaccessible to various factors and transcription proteins. Conversely, the formation of open and relaxed regions within the chromatin allows for pro-transcriptional machinery to bind and drive gene expression. Therefore, profiling chromatin accessibility can provide invaluable information regarding active regulatory regions and cell state. Short-read sequencing-based technologies have been widely used such as ATAC-seq, DNase-seq, and FAIRE-seq to profile chromatin accessibility [1–3]. Importantly, these short-read based methods lack the resolution to determine difference in accessibility between haplotypes. Recently, methods leveraging third generation long-read sequencing have also been developed. Long-reads provide more robust read mapping at repetitive sites and are suitable for haplotype-specific profiling. These methods can also identify DNA modifications. Therefore, such methods can leverage the use of DNA modifying enzymes to simultaneously sequence DNA and detect relevant modifications. Wang *et al* developed MeSMLR-seq by leveraging cytosine methylation using M.CviPI GpC methyltransferase and profiled accessible regions in yeast by relying the fact that GpC methylation does not present endogenously and M.CviPI can only affect DNA where it is accessible [4]. Lee *et al* also used M.CviPI and developed nanoNOMe to profile chromatin accessibility and DNA methylation in humans simultaneously [5]. Recently, Xie *et al* used M.CviPI treatment followed by DNA fragmentation and ligation and developed SCA-seq to profile spatial interaction of chromatin accessibility and DNA methylation [6]. GpC is not evenly distributed across the genome and there are regions with low GpC density. Moreover, GpC methylation can be confused with natural DNA CpG methylation in regions such as GCG. Shipony *et al* developed SMAC-seq by using EcoGII to convert adenine to N6-methyladenosine (6mA) and profiled accessible DNA using nanopore sequencing in yeast which lacks human genome complexity and endogenous DNA modifications [7]. The approach was unable to demonstrate accurate genome-wide chromatin accessibility in the human genome due to the lack of a high accuracy 6mA base calling model resulting in a high false positive rate. In addition to nanopore sequencing, other approaches utilizing Pacific Biosciences sequencing using 6mA have also been developed to profile chromatin accessibility [8,9].

Here we used EcoGII to convert adenine to 6mA and developed appropriate methodology and models for base calling and 6mA calling from nanopore data. Using these models, we then accurately profiled DNA methylation and accessibility in humans at haplotype level.

## Results and Discussion

To develop our models for base calling and 6mA detection of EcoGII treated samples, we first created a training dataset using the HG002 human sample. Purified DNA was treated with the non-specific 6mA transferase EcoGII to create a highly 6mA converted DNA sample and sequenced with the Oxford Nanopore (ONT) long-read platform. We assumed all adenines in this treated sample were converted to 6mA and used this sample along with sequence data from a HG002 untreated DNA sample as the training set [10]. To enable more robust base calling of reads containing 6mA, we first trained a base calling model and then we trained a model for 6mA calling. Our resulting base calling model outperformed ONT models with higher accuracy (ACC) and sequence identity (SI) (ACC= 92.9%; SI= 97.7%. For performance of ONT models see methods. Supplementary Figure 1). Our 6mA calling model also considerably outperformed ONT modification calling model (At 0.5 prediction threshold, Accuracy= 99.5%; Precision= 99.5%; Recall= 99.5%. See methods section for performance of ONT model. Supplementary Figure 1). We should point out that 6mA calling metrics are conservative estimates as we assumed that all the adenines in the EcoGII treated sample were converted 6mA which is unlikely to be true. However, there is no evidence regarding the presence of endogenous 6mA in human genome and estimates of false positive rate should hold true when evaluating unmodified DNA sequence [11]. With respect to the false positive rate (FPR), at a 0.5 prediction score threshold our method demonstrated 0.55% FPR while the ONT model had 33.81% FPR (Supplementary Figure 1). Additionally, we also used a MinION flow cell (FAB45280) public data from NA12878 to further examine FPR [12]. The Nanopore model demonstrated much better performance with 8% FPR compared to PromoethION data likely because the approach was initially developed using MinION data, however our model outperformed ONT model with a 0.44% FPR. Overall, both our base calling and 6mA calling models demonstrated robust performance and exceled ONT models (Supplementary Figure 1).

To demonstrate the ability of our model to profile 6mA we used the NA12878 and MCF7 cell lines. Prior to long read sequencing we treated NA12878 and MCF7 chromatin samples with EcoGII. Where accessible DNA regions should be available to this enzyme while inaccessible regions should not. We then sequenced these samples and called DNA 6mA using the trained model. NA12878 was sequenced to redundant fold coverage of ∼25 and MCF7 to ∼14. We also called 5mC for these samples using nanopolish [13]. The 5mC data demonstrated a high correlation with the publicly available whole genome bisulfite sequencing (WGBS) data for NA12878 and MCF7 [14,15], with a Pearson correlation of 0.895 and 0.837, respectively (Figure 1a). To further demonstrate the utility of our approach, we profiled DNA methylation and accessibility at CpG islands (CGIs). CpG islands usually lack CpG methylation and are accessible and so should also correlate with higher levels of 6mA conversion. As demonstrated in Figure 1b and 1c, CpG methylation is low while DNA accessibility (6mA) is high at CGIs.

DNA methylation and accessibility profiles from our approach closely follows the trend from other methods (WGBS for CpG methylation and ATAC-seq, DNase-seq and FAIRE-seq for chromatin accessibility for these cell-lines). We profiled DNA methylation and accessibility with respect to the transcription start site (TSS) of silenced and highly expressed genes (Figure 1d and 1e). Using public RNA-seq data we determined silenced genes (transcript per million or TPM=0) and the top ten thousand highly expressed genes in NA12878 and MCF7 [14]. Our DNA methylation and accessibility profiles agreed with other methods and highly expressed genes demonstrated low CpG methylation and high accessibility while silenced genes showed high methylation and low accessibility. As for CGIs, DNA accessibility and methylation profiles from our approach closely replicated the observation from other methods (Figure 1d and 1e).

**Figure 1.**
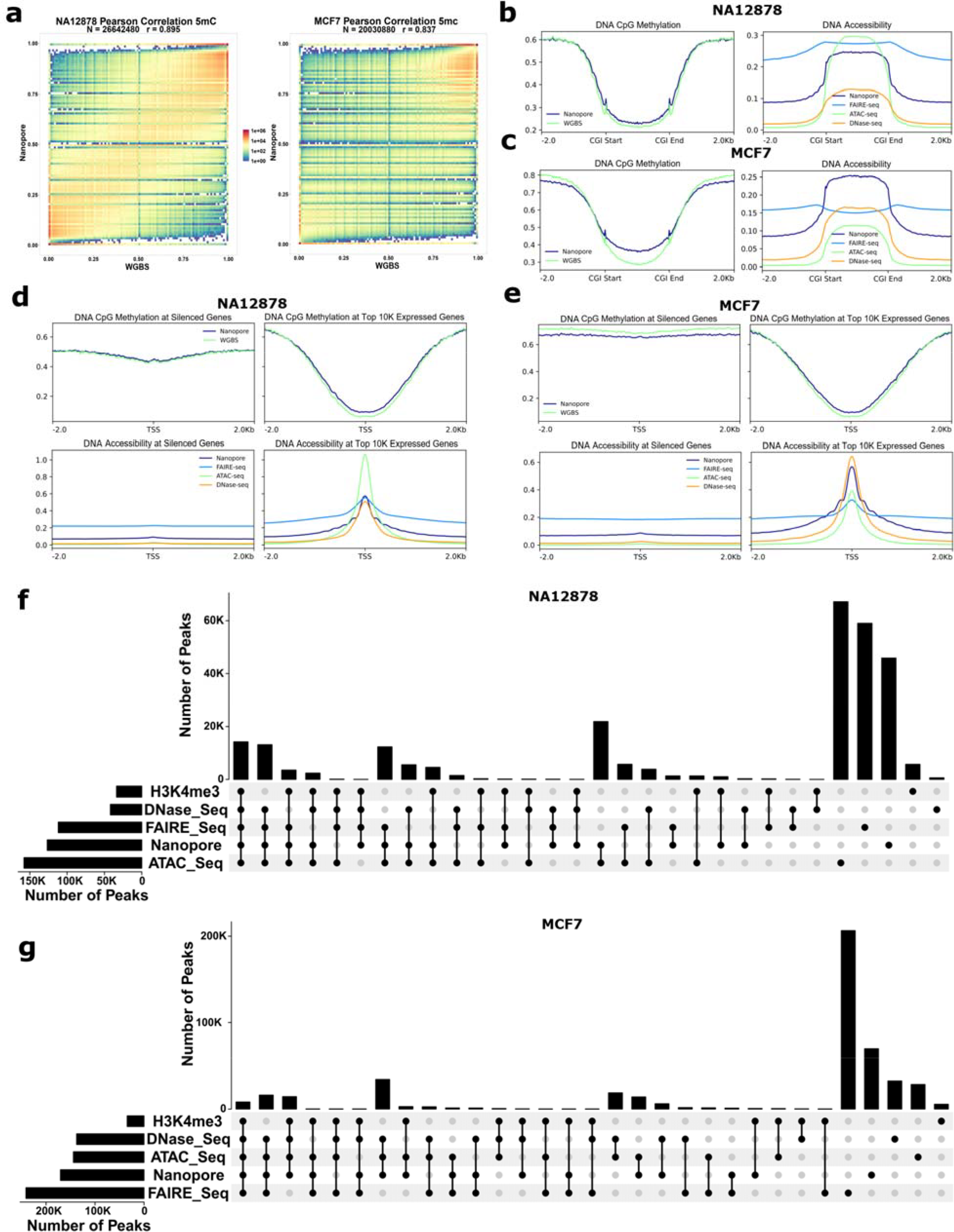
Simultaneous profiling of DNA methylation and accessibility. a) Pearson correlation of CpG methylation frequencies between nanopore and WGBS in NA12878 and MCF7 cell lines. CpG sites with a coverage >4 was used for correlation analysis. b,c) Metaplots of DNA methylation and accessibility at CpG islands. d,e) Metaplots of DNA methylation and accessibility around transcription start site (TSS) at the top 10K highly expressed genes and silenced genes (TPM=0) in NA12878 and MCF7 cell lines. For the sake of visualization, all the DNA accessibility signals from other approaches were scaled to be close to 6mA frequency (0-1) from nanopore. FAIRE-seq signals multiplied to 50. DNase signals multiplied to 0.25. ATAC-seq signals multiplied to 0.001. f,g) Overlapping DNA accessibility peaks detected using nanopore 6mA to peaks from other methods.

Next, we sought to find peaks based on 6mA frequency results that demonstrate accessible regions across the genome. For this, we performed differential 6mA methylation analysis and detected differentially accessible regions (DARs) showing higher 6mA compared to the background untreated HG002 sample which should lack 6mA (Methods). Such DARs are basically demonstrating accessible regions in the chromatin. Using this approach, we detected ∼141K peaks in NA12878 and ∼182K peaks in MCF7 (Supplementary Tables 1 and 2). For comparison, we used chromatin accessibility peaks from ATAC-seq, DNase-seq, and FAIRE-seq. Because H3K4me3 is a histone mark of accessible/active regions, we also used H3K4me3 ChIP-seq peaks [16]. We gathered ∼229K peaks for NA12878 and ∼403K peaks for MCF7 from public ATAC-seq, DNase-seq, FAIRE-seq, and H3K4me3 ChIP-seq data [14]. In NA12878, 68% of the nanopore peaks overlapped to 36% of all the peaks (85% DNase-seq, 79% H3K4me3, 51% ATAC-seq, 42% FAIRE-seq) gathered from public data (Figure 1f and Figure 2a). In MCF7, 62% of nanopore peaks overlapped to 26% of all the peaks (80% H3K4me3, 64% ATAC-seq, 61% DNase-seq, 12% FAIRE-seq) gathered from public data (Figure 1g and Figure 2a). Looking into overlaps of the peaks from different methods, nanopore peaks demonstrated robust overlap to gold standard ATAC-seq and DNase-seq. As demonstrated in Figure 1f and 1g, all methods of chromatin accessibility demonstrated comparable overlap and non-overlap peaks, however in the MCF7 sample FAIRE-seq seems to be an outlier with many peaks unique to this approach.

**Figure 2.**
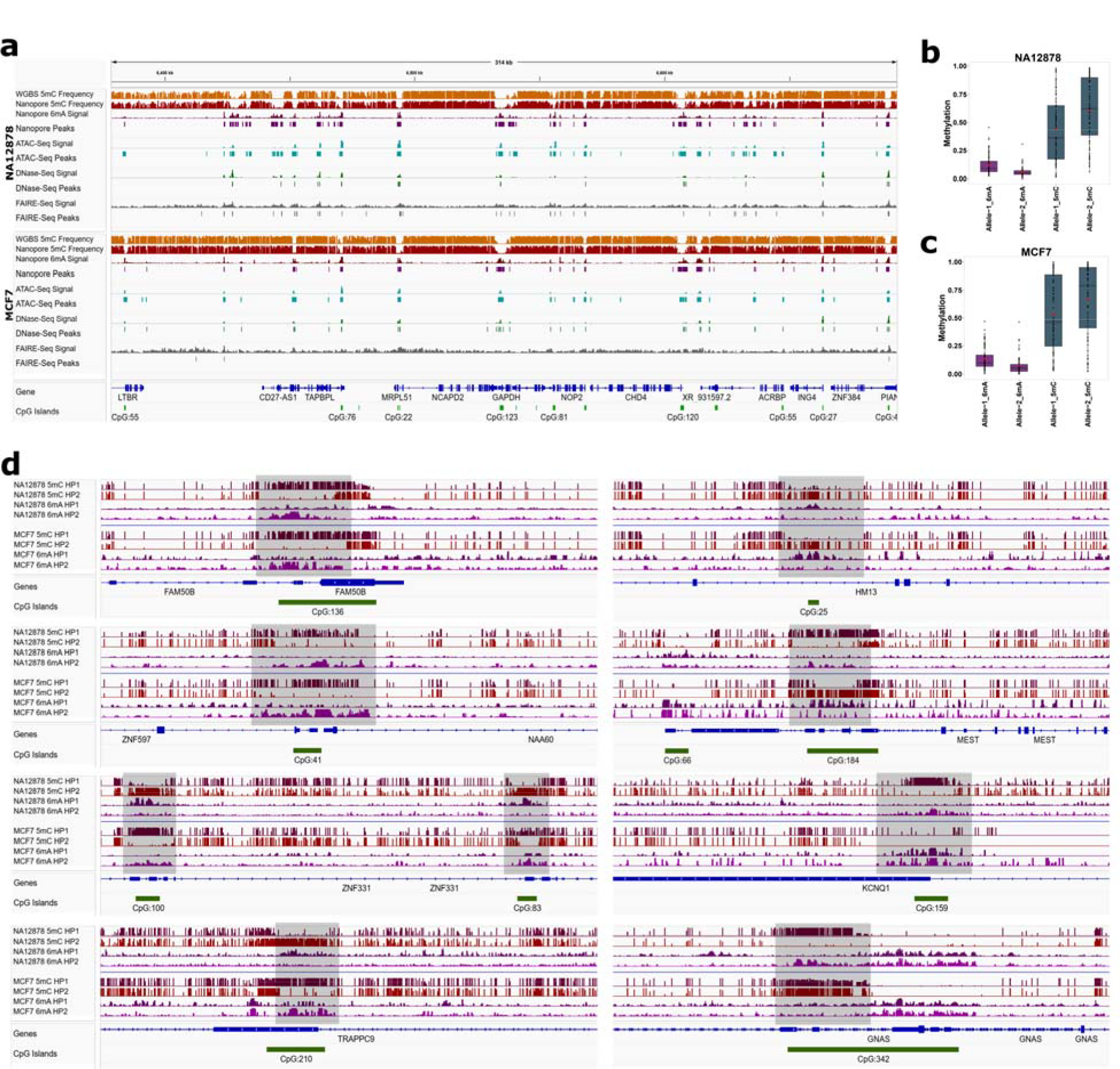
DNA methylation and accessibility tracks and haplotype specific results. a) A 314Kb interval from IGV that demonstrated DNA accessibility signals and peaks using nanopore and other approaches. b,c) Average of adenine and CpG methylation for each allele at well-known imprinted regions. Hypomethylated alleles show higher 6mA (DNA accessibility) while hypermethylated alleles show lower 6mA. For the sake of visualization, haplotype 1 and haplotype 2 were swapped so that haplotype 1 (or allele 1) always have the higher 6mA at each iDMR. If a haplotype was swapped for 6mA it was also swapped for 5mC. d) IGV screenshots at several known imprinted regions demonstrate allele specific CpG methylation and 6mA. HP1 represents haplotype 1 and HP2 shows haplotype 2.

A key advantage of using long-read sequencing is the ability to haplotype reads and detect allele-specific events. We therefore phased nanopore reads and 6mA and CpG methylation (Methods) and investigated allele-specific DNA methylation and accessibility at known imprinted regions where it is known that only one allele is methylated and the other allele is not methylated and is accessible to regulatory proteins. We used the well-known imprinted differentially methylated regions (iDMRs) we previously gathered [10], that was reported by at least two studies [17–21]. In general, 6mA and 5mC methylation demonstrated opposite direction at the iDMRs where allele with high CpG methylation showed low 6mA and vice versa (Figure 2b-d). In some iDMRs (e.g. *KCNQ1/KCNQ1OT1* and *GNAS-Ex1A*) allelic methylation and accessibility is present in one cell line and not the other, possibly due to the tissue variation or polymorphic imprinting or due to loss of imprinting in cancer.

To further analyse allele-specific events we focused on NA12878 which is a normal (i.e., non-tumour derived) female cell line. We called allele specific differentially accessible regions (aDARs) and differentially methylated regions (aDMRs) by performing differential 6mA and CpG methylation analysis between haplotypes (Methods). We detected 7,932 aDMRs and 11,791 aDARs in NA12878 (Supplementary Tables 3 and 4). As observed previously [5], the overlap between allelic DARs and DMRs was small (427 aDARs), however the majority of overlaps (412) were concordant where the CpG methylated allele had low 6mA/accessibility and vice versa. 69.9% (65 iDMRs) and 23.7% (22 iDMRs) of the well-known iDMRs overlapped with aDMRs and aDARs, respectively (Supplementary Tables 5 and 6). We mapped aDARs and aDMRs to the chromosome X inactive and escapee genes we previously gathered [22]. Inactive genes are only expressed from one allele while escapee genes are biallelically expressed, and we expect inactive genes demonstrate aDAR and aDMR at promoter. As expected, aDARs and aDMRs mapped to the promoter of inactive genes in a greater proportion (Supplementary Tables 7 and 8). Allelic DMRs mapped to the promoter (1.5Kbp upstream and 500bp downstream of transcription start site) of 41.2% (153 genes) of X-inactivated and 16% (12 genes) of X-escapee genes (Supplementary Table 7). Allelic DARs mapped to the promoter of 21.8% (81 genes) X-inactivated and 2.7% (2 genes) of X-escapee genes (Supplementary Table 8). Active genes tend to be CpG methylated in their gene bodies and gene-body methylation on the active X-chromosome genes compared to the inactive X-chromosome has been observed [23,24]. Therefore, we mapped aDMRs to the gene body. 64 genes (61 X-inactivated and 3 X-escapee) had aDMR at the promoter and body and most of the aDMRs demonstrated the expected methylation direction where methylated allele at the promoter demonstrated lower methylation at the gene body and vice-versa (Supplementary Table 7). On the other hand, aDARs demonstrate higher accessibility on active X-chromosome at different genomic context such as gene body and promoter [5]. 29 genes (27 X-inactivated and 2 X-escapee) had aDAR at the promoter and body and the majority of the aDARs on the gene body and promoter showed the same direction where allele with higher accessibility at gene promoter was also more accessible at the gene body (Supplementary Table 8).

## Conclusions

In summary, our base calling and 6mA calling models demonstrated robust performance and enabled simultaneous profiling of DNA accessibility and methylation in human with high resolution with only ∼25 fold coverage in NA12878 and ∼14 fold coverage in a MCF7 cell line sample. Additionally, the protocol involves a relatively simple enzymatic treatment of chromatin before library construction. We also showed the ability of our approach to profile haplotype-specific features. Therefore, our approach demonstrates a methodology for high resolution profiling of DNA accessibility and CpG methylation in complex mammalian genome.

## Methods

### HG002, NA12878, and MCF7 cell culture and EcoGII treatment

HG002 (GM24385) cells were cultured in RPMI1640 with 15% fetal bovine serum (not heat inactivated). NA12878 human lymphoblastoid cells were maintained in RPMI Medium 1640 supplemented with 2mM L-glutamine and 10% fetal bovine serum. MCF-7 human breast cancer cells were grown in DMEM supplemented 10% FBS and the cells were kept at 37 °C with 5% CO_2_.

EcoGII Methyltransferase is a non-specific methyltransferase that modifies adenine residues (N6). For HG002 treatment, we generated the high-molecular-weight DNA from HG002, using Magattract kit (67563, Qiagen). The input DNA was 10ug (75ng/ul) and the reaction was setup as follows according to the manufacturer’s instructions, with slight modifications: 10xCutSmart buffer 50μl, diluted 1600uM S-Adenosyl methionine (SAM; prepared from 32mM Stock) 50ul, 200U EcoGII Methyltransferase enzyme (New England BioLabs, M0603S), Nuclease-free Water up to 500μl. The reaction was incubated at 37°C for 6 hours in a Thermomixer. Additional 160 µM S-Adenosyl methionine (SAM) was spiked in after 2 hours and 4 hours respectively. After 6 hours of total incubation, the reaction was stopped by heating at 65°C for 20 minutes, followed by sodium acetate and ethanol precipitation and sequencing. For NA12878 and MCF-7 treatment, intact chromatin was treated with EcoGII Methyltransferase as previously suggested with minor modifications [7]. Briefly, 5 × 10^6^ MCF-7 and NA12878 cells were washed with 1× PBS, then resuspended in 200 μl of ice-cold nuclei lysis buffer (10 mM Tris pH 7.4, 10 mM NaCl, 3 mM MgCl2, 0.1 mM EDTA, 0.5% NP-40) and incubated on ice for 10 min. Nuclei were then centrifuged at 500g for 5 min at 4°C, resuspended in 200 μl of cold nuclei wash buffer (10 mM Tris pH 7.4, 10 mM NaCl, 3 mM MgCl2, 0.1 mM EDTA) and centrifuged again at 500g for 5 min at 4 °C. Finally, nuclei from 2 million MCF-7/NA12878 cells at a viability of ∼95% were resuspended in 200 μl of reaction buffer (1× NEB CutSmart buffer, 0.3 M sucrose). Nuclei were then treated with EcoGII by adding 200 U of EcoGII (NEB) and SAM at 0.6mM and incubated at 37°C for 1hr with gentle shaking at 900rpm. The reaction was stopped by adding 0.2% SDS, followed by HMW DNA extraction using Magattract kit (67563, Qiagen) followed by sequencing.

### Library construction and sequencing

Sequencing library construction and sequencing were performed as per manufacturer’s protocol SQK-LSK110 (Oxford Nanopore Technologies). Libraries each with 2ug of input DNA were prepared, conforming to Oxford Nanopore Technologies’ protocols, using the SQK-LSK110 Ligation Library Kit. No DNA shearing was performed. The NEB UltraII kit (New England Biolabs, Ipswich, MA, USA, cat. no. E7646A) was used for end-repair and A-tailing. NEBNext quick ligase (E6056S) was used to ligate the Oxford Nanopore sequencing adapter. A final size selection of 0.4:1 ratio (magnetic beads to the library) was done to select against smaller molecules. Automated library prep was done using the Hamilton Nimbus liquid handling platform.

DNA libraries were loaded in R9.4.1 pore flow cells on PromethION 24 instrument running software version 19.06.9 (MinKNOW GUI v4.0.23). Sequencing was carried out for 72 hours. DNase I (Invitrogen cat no. AM2222) nuclease flush was performed after 24-48 hours by reloading the flow cell with the same library mix.

### Model training

We trained a base calling model and a 6mA calling model using our EcoGII treated HG002 sample and an untreated HG002 sample from our previous study [10]. To train the base calling model, we used the ONT Taiyaki tool (https://github.com/nanoporetech/taiyaki). First, signals from both samples were mapped to the reference genome (hg38_no_alt) using ONT megalodon v2.5.0 tool (https://github.com/nanoporetech/megalodon) with guppy server v6.1.1 and *dna_r9*.*4*.*1_450bps_sup_prom* config and model. Mapped signals from both samples were then merged using *merge_mappedsignalfiles*.*py* script from taiyaki with *--batch_format* option selected. Merged signal file was then used to train the base calling model using *train_flipflop*.*py* script from taiyaki with *mLstm_flipflop*.*py* model.

To train the 6mA calling model, we used ONT Remora v1.0.0 (https://github.com/nanoporetech/remora). First signals from treated and untreated samples were mapped to the reference genome (hg38_no_alt) using ONT megalodon v2.5.0 with guppy server v6.1.1 and the base calling model we trained with *dna_r9*.*4*.*1_450bps_sup_prom* configuration (For the EcoGII treated sample *--ref-mods-all-motifs* option was used to allow modification mapping for signals). Mapped signals from both samples were then merged using *merge_mappedsignalfiles*.*py* script from taiyaki with *--allow_mod_merge* and *--batch_format* options selected. Merged signal file was then used to train the 6mA calling model using remora as follow: First, *remora dataset prepare* with options *--motif A 0 --chunk-context 250 250 --kmer-context-bases 9 9 --max-chunks-per-read 15* command was used to prepare the mapped signals for training. Afterwards, the prepared training dataset was used to train the 6mA calling model using *remora model train* command with the setting *--model ConvLSTM_w_ref*.*py --size 128 --balance --epochs 25*. To test the model, we used 135K unseen reads from untreated/native HG002 sample and 300K reads from EcoGII HG002 treated sample and we considered all the adenines of untreated sample as canonical A and all the adenines from treated sample as 6mA. Our base calling model outperformed ONT models with considerably higher accuracy (ACC) and sequence identity (SI) in the treated sample (Our model: ACC= 92.9%; SI= 97.7%. ONT modification model: ACC= 78.1%; SI= 89.9%. ONT super accuracy model: ACC= 77.8%; SI= 88.1%). In the untreated sample our model outperformed ONT modification model but showed slightly lower accuracy and sequence identity compared to ONT super accuracy base calling model (Our model: ACC= 92%; SI= 97.3%. ONT all context model: ACC= 89.5%; SI= 96.3%. ONT super accuracy model: ACC= 93.9%; SI= 97.7%). Our 6mA calling model considerably outperformed ONT modification calling model (At 0.5 prediction threshold, our model: Accuracy= 99.5%; Precision= 99.5%; Recall= 99.5%. ONT model: Accuracy= 84.5%; Precision= 75.9%; Recall=99%. (Supplementary Figure 1).

### 6mA calling for MCF7 and NA12878 chromatin EcoGII treated and untreated HG002 samples

Megalodon v2.5.0 with guppy server v6.2.1 and the base calling (with *dna_r9*.*4*.*1_450bps_sup_prom* configuration) and 6mA calling models we trained with *--mod-binary-threshold 0*.*75* was used to baseball and 6mA call nanopore data. Base called reads were aligned to the human reference genome (hg38_no_alt) using minimap2 v2.24 and variants were called using clair3 v0.1-r10 with *r941_prom_sup_g5014* model [25,26]. Clair3 called variants were then phased using WhatsHap v1.2.1 [27]. To detect DNA accessibility peaks we used the DSS R package v2.46.0 and performed differential 6mA analysis against untreated HG002 sample to detect DMRs/peaks [28]. We only used peaks with >0.15 adenine methylation compared to untreated sample.

To detect allele specific 6mA, we haplotagged the bam file using WhatsHap and then extracted haplotype 1 and 2 reads, we then used *megalodon_extras aggregate* command with haplotype 1 or haplotype 2 reads for the *--read-ids-filename* option with *--mod-binary-threshold 0*.*75* to profile allele-specific 6mA. To call aDARs, we used DSS R package v2.46.0 and performed differential 6mA analysis between haplotypes [28]. We only used aDARs with |Methylation difference| >0.2.

### 5mC calling for MCF7 and NA12878 chromatin EcoGII treated samples

For CpG methylation calling, we used nanopolish v13.3 software and used *--call-threshold 1*.*5* to obtain per-site CpG methylation frequencies [13]. After CpG methylation calling, we used NanoMethPhase v1.2.0 to detect allele specific CpG methylation [22]. We then detected aDMRs using NanoMethPhase *dma* module with DSS v2.46.0. We only considered aDMRs with |Methylation difference| >0.2.

### Public data for comparison

We used public WGBS data for comparison to our CpG methylation results and ATAC-seq, DNase-seq, and FAIRE-seq peak calling and signal data for comparison to our chromatin accessibility results.

WGBS CpG methylation data obtained from gene expression omnibus for NA12878 (GSM2308633) and MCF7 (GSM3336908) [14,15]. Strand-level CpG methylation data were aggregated (# of reads called methylated from both strands / # of all reads from both strands) to make a consensus methylation frequency for each CpG site. Chromatin accessibility peaks and signals obtained from ENCODE [14]. ATAC-seq (ENCFF285BIH and ENCFF667MDI), DNase-seq (ENCFF412PRG and ENCFF960FMM), and FAIRE-seq (ENCFF000THW and ENCFF000THZ) obtained for NA12878. ATAC-seq (ENCFF502RTI and ENCFF976UNK), DNase-seq (ENCFF609LOE and ENCFF134COA), and FAIRE-seq (ENCFF000TMT and ENCFF000TMW) obtained for MCF7 cell line. Overlapping peaks in each dataset were merged. FAIRE-seq results were based on hg19. Therefore, we lifted over hg19 data to hg38 using Crossmap.py and hg19Tohg38 chain file from UCSC genome browser [29,30]. To find silenced and highly expressed genes, gene expression data for NA12878 (ENCFF545OJE) and MCF7 (ENCFF395RXK) was also obtained from ENCODE [14]. Top 10K genes with highest TPM were used as highly expressed genes and all the genes with TPM of 0 were used as silenced genes.

## Availability of data and materials

Trained models and codes are provided at GitHub (https://github.com/vahidAK/6mA_Work). CpG methylation (5mC) and 6mA frequency data for NA12878 and MCF7 EcoGII-treated chromatin samples are available from Zenodo (DOI: https://doi.org/10.5281/zenodo.8407960). 6mA frequency data for native (untreated) HG002 sample is available from Zenodo (DOI: https://doi.org/10.5281/zenodo.8408392). Raw and base called nanopore sequencing data will be available through the Sequence Read Archive (SRA).

## Authors’ contributions

All authors jointly conceived the project. V. A developed the analysis pipeline and performed dry-lab experiments and analysis including model training, base calling, 6mA and 5mC calling, and comparisons to other methods. S. L performed wet-lab experiment including cell cultures, DNA and chromatin extractions and enzymatic treatments. V. A wrote the whole manuscript with input regarding the cell culture and sequencing in the “Methods” from S. L. SJMJ supervised the project and acquired funding. All authors reviewed and edited the manuscript.

## Acknowledgements

We appreciate contributions from the Library Construction and Sequencing teams at BC Cancer Genome Sciences Centre. VA acknowledges funding from the University of British Columbia Four-Year Doctoral Fellowship. SJMJ acknowledges funding from the Canada Research Chair program.

