## Supplementary Figure 1 for "Profiling Chromatin Accessibility in Humans Using Adenine Methylation and Long-Read Sequencing"

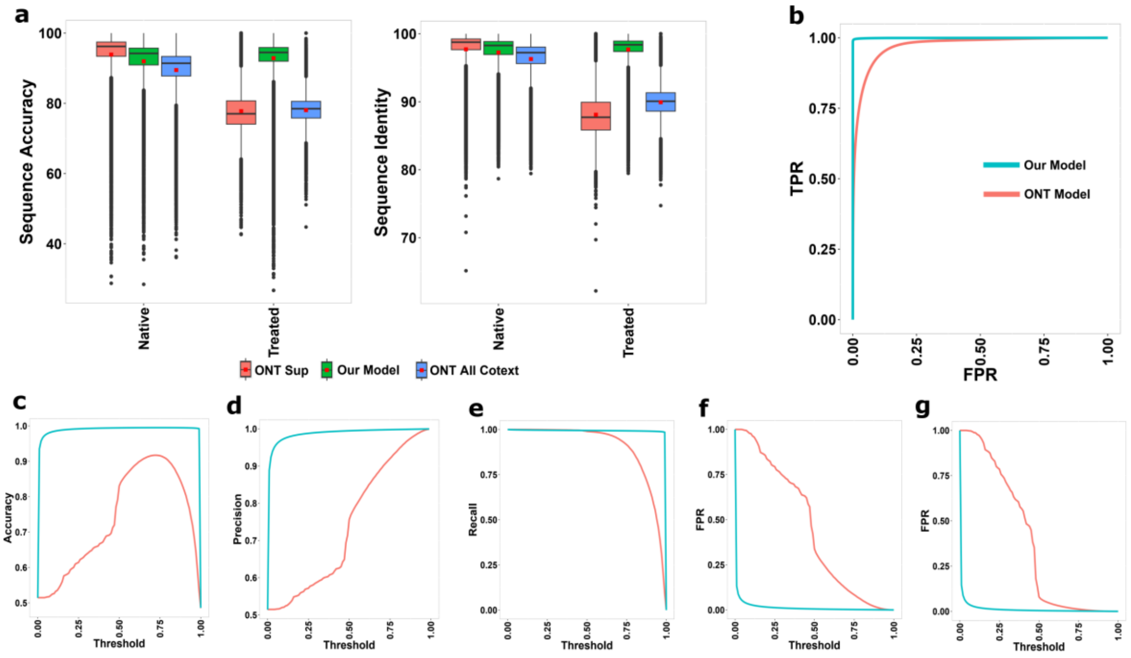


Supplementary Figure 1. a) Read level base calling performance of our base calling model compared to ONT modification and super accuracy models. b) Receiver operating characteristic (ROC) curve of 6mA calling using our model and ONT all-context model. c,d,e) Accuracy, precision, and recall plots at various prediction thresholds for our 6mA calling model and ONT all-context model. f) False positive rate (FPR) for our 6mA model and ONT all-context model at various prediction thresholds. In a-f, comparisons are based on 300K unseen reads from HG002 treated sample and 135K unseen reads from HG002 native/untreated sample. All adenines from treated sample considered as 6mA and all adenines from untreated sample considered as A. g) FPR for our 6mA model and ONT all-context model at various prediction thresholds based on ~128K reads from NA12878 MinION sample.
